# Use-Dependent Plasticity Enhances Nociceptive and Non-Nociceptive Somatosensory Cortical Responses in Healthy Individuals

**DOI:** 10.1101/2020.10.29.360222

**Authors:** A.M. Zamorano, B. Kleber, F. Arguissain, P. Vuust, H. Flor, T. Graven-Nielsen

## Abstract

Repetitive movements and multisensory integration promote use-dependent plasticity, which has been consistently demonstrated using musical training as a model. Prolonged and repeated execution of motor patterns is also a main risk factor for developing pain syndromes, yet the neural underpinnings that link repetitive movements and use-dependent plasticity to abnormal pain processing in humans are unknown. Employing nociceptive and non-nociceptive electrical stimulation to evoke brain responses, we demonstrate that healthy musicians compared to non-musicians show larger non-nociceptive N140 and nociceptive N200 responses, smaller non-nociceptive P200 responses, and faster reaction times (RTs) to both stimuli. Across participants, larger N140 and N200 amplitudes predicted the RTs, whereas amount of daily practice in musicians predicted RTs, non-nociceptive P200, and nociceptive P300 components. These novel findings indicate that the mechanisms by which repetitive sensorimotor training and multimodal integration promote neural plasticity in multisensory neural structures may also increase stimulus-selectivity to nociceptive cues in healthy individuals.

## INTRODUCTION

Repetitive movements and their associated multisensory integration play a major role in the structural and functional reorganization of sensory and motor neural connections (Bütefisch et al., 2000; Classen et al., 1998; Recanzone et al., 1993; Zatorre et al., 2012). This phenomenon, known as use-dependent plasticity, has been widely investigated in experienced musicians, as musical training is a popular *in vivo* model for evaluating the effects of extensive sensorimotor training (Herholz and Zatorre, 2012; Jäncke, 2009). Musicians typically begin with deliberate daily practice routines in early childhood and develop expert perceptual and motor skills during adolescence and adulthood, which in turn have been associated with a number of functional and structural adaptive changes in brain regions involved in sensory perception (Kraus and Chandrasekaran, 2010; Pantev et al., 1998), sensorimotor control (Kleber et al., 2013; Rosenkranz et al., 2007), and cognitive functions (Brown et al., 2015). Despite such adaptive effects, animal models have shown that extensive repetitive movements and associated sensory integration (i.e. sensorimotor training) can also contribute to the genesis of maladaptive neural plasticity assumed to be involved in the development of pain syndromes and focal dystonia (Byl et al., 1996). In humans, the prolonged and repeated execution of motor patterns is also considered one of the main risk factors for developing pain musculoskeletal syndromes (Herin et al., 2014), yet the neural underpinnings that link repetitive movements to abnormal pain processing are still unknown.

Interestingly, musculoskeletal pain syndromes and focal dystonia are significantly more prevalent in musicians than in the general population (Rotter et al., 2020; Sadnicka et al., 2018). In line with these observations, a behavioral study with healthy musicians provided first evidence suggesting that the explicit training of perceptual and motor skills may not only be associated with enhanced musically relevant functions but also with increased pain sensitivity (Zamorano et al., 2015). In addition, recent resting-state fMRI studies revealed enhanced insula-based connectivity in pain-free musicians, comparable to changes observed in chronic pain non-musicians (Zamorano et al., 2019, 2017), indicating that extensive sensorimotor training and chronic pain may affect overlapping neural systems. In order to test this assumption, the current study directly explored the nociceptive and non-nociceptive somatosensory pathways to determine the neural foundation by which extensive sensorimotor training may alter nociceptive processing in healthy individuals.

The underlying hypothesis is that the same processes by which extensive sensorimotor training and multisensory integration can modify task-specific topographic and functional neural representations in the brain (Byl et al., 1996; Elbert et al., 1995), as well as facilitate the priming of neural responses in brain areas where the processing of multimodal stimuli converges (Paraskevopoulos et al., 2012), may also shape nociceptive neural and behavioral responses, as previously indicated by invertebrate models (Hu et al., 2017; Ohyama et al., 2015). Nociceptive inputs are conveyed via the spinothalamic pathways to the brain, engaging multiple regions such as the primary and secondary somatosensory cortices, the prefrontal cortex, the insula, and the anterior cingulate cortex, as well as subcortical areas like the thalamus (Apkarian et al., 2005). The sensory cortices, the insula, and the cingulate cortex are also commonly activated during musical temporal processing (Platel et al., 1997), sound integration (Bamiou et al., 2003), the acquisition of action–perception links (Mutschler et al., 2007), and sensorimotor control (Kleber et al., 2017). Likewise, non-nociceptive inputs, which commonly convey via the dorsal column–lemniscal pathways, can also converge in the nociceptive pathways by gaining access to wide dynamic range (WDR) neurons in the spinal cord (D’Mello and Dickenson, 2008). Based on this notion and our observations of increased pain sensitivity and insula connectivity in healthy musicians (Zamorano et al., 2019, 2017, 2015), we propose that repetitive sensorimotor training and multimodal integration can also prime the insular and cingulate responses to other sensory processes, such as nociception.

To test this assumption, we used nociceptive and non-nociceptive electrical stimulation paradigms to assess the spinothalamic and the dorsal column–lemniscal pathways with the aim of testing whether extensive sensorimotor training can facilitate the transient brain responses in healthy individuals. By analyzing the neural and behavioral response components as a function of accumulated sensorimotor training (i.e., musical practice), we also aimed to characterize how extensive multisensory training may influence the variability of cortical responses to noxious and non-noxious stimuli across individuals. Following our hypotheses, we expected enhanced nociceptive and non-nociceptive evoked responses in healthy musicians relative to non-musicians. Moreover, we expected that the amount of sensorimotor musical training would predict the variation of the evoked brain responses and reaction times.

## MATERIALS AND METHODS

### Participants

Previous electrophysiological studies exploring use-dependent plasticity are typically based on a total sample size between 15-44 participants (Kühnis et al., 2014; Pantev et al., 2001; Vuust et al., 2008). Moreover, studies using the same intra-epidermal electrical stimulation to elicit nociceptive electrophysiological responses as in the current study report total sample sizes of 5-20 participants (Biurrun Manresa et al., 2018; Mouraux et al., 2014, 2010; van den Berg et al., 2020). To ensure a robust estimate of our neurophysiological effect, a total sample of 40 participants was recruited for the current study. Twenty trained musicians (nine female, 18 right-handed, mean age 26.5 ± 3.8 years,) participated in this study. All musicians were conservatory-trained instrumentalists (6 string, 3 keyboard, 8 woodwind, and 3 brass instruments) with a long history of professional experience, including a total average of 18,102 hours (± 8,322 hours) of musical practice and a daily average of 4.2 hours (± 2 hours). The control group included 20 non-musicians (nine female, 19 right-handed, mean age 26.9 ± 5.3 years) without any prior formal or informal music training. Exclusion criteria were neurological, cardiorespiratory, or mental disorders, as well as chronic pain, or pregnancy. All participants received written and verbal information about the scope of this study and provided written consent. The study was performed in accordance with the Declaration of Helsinki (General Assembly of the World Medical Association, 2013) and approved by the local ethics committee (Den Videnskabsetiske Komité for Region Nordjylland, N-20170040).

### Experimental procedure

Participants were seated in a comfortable chair and familiarized with the electrical test stimuli. To avoid excessive head and body movements, participants were instructed to fixate their gaze on a black cross (3 x 3 cm) displayed 1.5 m in front of them for the entire duration of each stimulation block.

The experiment consisted of two stimulation blocks with a sequence randomized and counterbalanced across participants. Each block was comprised of 30 stimuli belonging to one of two types of electrical stimulation to the right hand: (1) intra-epidermal electrical stimulation, which predominantly activates Aδ nociceptors (Mouraux et al., 2010), and (2) low-intensity transcutaneous electrical stimulation, which activates non-nociceptive Aβ fibers (Burgess and Perl, 1967). To ensure that each stimulus was perceived and to maintain vigilance across time, participants had to press a button immediately after the perception of each stimulus (reaction time). Detection thresholds were recorded for each stimulation modality at baseline.

### Electrical stimulation

To ensure that the stimuli remained selective for their respective fibers, the intensity was individually adjusted to twice the detection threshold (Mouraux et al., 2010). Both nociceptive and non-nociceptive stimuli consisted of two rapidly succeeding constant-current square-wave pulses with a duration of 0.5 ms each, an inter-pulse interval of 5 ms, and an inter-stimulus interval that randomly varied between 8 and 10 s (Mouraux et al., 2014). The electrical stimuli were controlled using custom-made software (“Mr. Kick”, Aalborg University, Aalborg, Denmark), and delivered by a constant-current electrical stimulator (Digitimer DS5, Digitimer Ltd., Welwyn Garden City, UK).

Nociceptive stimuli were delivered using intra-epidermal electrical stimulation (Inui et al., 2002). Stimuli were delivered using a stainless steel concentric bipolar needle electrode developed by Inui et al. (2002), consisting of a needle cathode (length: 0.1 mm, Ø: 0.2 mm) surrounded by a cylindrical anode (Ø: 1.4 mm). Gently pressing the device against the skin inserted the needle electrode into the epidermis of the dorsum of the right hand, which clearly elicited a burning/pricking sensation when stimulated. These stimuli, provided that low intensities are used, predominantly activate nociceptive Aδ fibers (Mouraux et al., 2010).

Non-nociceptive stimuli were elicited using low-intensity transcutaneous electrical stimulation. Stimuli were delivered through a pair of digital ring electrodes (Digitimer Ltd., Welwyn Garden City, UK) and applied to the first two phalanges of the right index finger (1-cm inter-electrode distance). These stimuli, provided that low intensities are used, predominantly activate non-nociceptive Aβ fibres (Burgess and Perl, 1967).

### Behavioral measures

Detection thresholds for nociceptive and non-nociceptive stimuli were estimated using a staircase procedure. The initial stimulus intensity was 30 μA for the nociceptive and 100 μA for the non-nociceptive stimulation, and the initial step sizes were 50 μA and 500 μA, respectively. After the first staircase reversal, step size was reduced to 10 μA and 100 μA, respectively. The procedure was interrupted after the occurrence of three staircase reversals at final step size. The detection thresholds were estimated by averaging the intensity of the stimuli at which these three reversals occurred.

The participants were instructed to push a button held in the left hand as soon as they perceived the stimulus. The mean reaction time (RT) across the 30 stimulations recorded relative to stimulus onset was extracted. RTs greater than 1000 ms were considered as undetected. The frequency distribution of RTs according to stimulus type was analyzed.

### Electrophysiological measures

Electroencephalographic (EEG) activity was recorded using an active electrode cap (g.SCARABEO, g.tec, Medical Engineering GMBH, Austria). The electrode montage included 64 electrodes according to the modified 10-20 system. During the recording, the EEG signals were amplified and digitized using a sampling rate of 1200 Hz and a left earlobe (A1) reference (g.Hlamp, g.tec, Medical Engineering GMBH, Austria). The ground electrode was placed at position AFz. The impedance of all electrodes was kept below 20 kΩ and assessed by the EEG system software (g.Recorder, g.tec, Medical Engineering GMBH, Austria).

Event-related potentials (ERPs) were analyzed offline using EEGLAB v.14.1.1 (Delorme and Makeig, 2004) running under MATLAB (The Mathworks, Natick, Massachusetts, USA). Data were band-pass filtered (0.5 - 40 Hz), followed by an Infomax independent component analysis using the in-built EEGLAB function runica to identify and remove components associated with noise (e.g., eye movement, eye blinks, cardiac, and muscle artefacts) (Jung et al., 2000). Continuous data were segmented into 1.5 s epochs, stimulus-locked from −500 to 1000 ms with time 0 corresponding to the stimulus onset. Baseline correction was made using the −500 to −10 ms window. For each subject and stimulus type, baseline-corrected epochs were further averaged to extract the ERPs of interest (Kunde and Treede, 1993; Mouraux et al., 2010).

For the ERPs in response to nociceptive stimuli, N100, N200, and P300 components were analyzed. The N100 component, commonly labelled in pain research as N1, was defined as the first major negative deflection occurring within the 60 ms time window preceding the N200 component (i.e., 100 ms - 160 ms), and measured with the recommended temporal–frontal montage (T7-Fz) (Valentini et al., 2012). The N200 and P300 components, labelled in pain research as N2 and P2, respectively (Cruccu et al., 2008), were identified with the recommended central-earlobe montage (Cz-A1). The N200 was defined as the first major negative deflection after stimulus onset, while P300 was defined as the first major positive deflection occurring after stimulus onset (Cruccu et al., 2008). For the ERPs in response to non-nociceptive stimulation, the N140 (analogous to N200) and P200 were determined using the midline Cz-A1 montage (Shimojo et al., 2000).

Exploratory statistical analyses were performed in the following ERP components to assess the effects of extensive sensorimotor training: non-nociceptive P50, P100, and P300 as well as a sustained contralateral negativity (labelled SCN) in the nociceptive stimulation. These peak latencies were chosen on the basis of previous research (Katus et al., 2015; Polich, 2007), as well as visual inspection of the grand averages.

The amplitudes of the ERP components were extracted from specific time windows of interest, which were centered at the peak latency of each ERP component and extended before and afterwards accordingly. The following time windows for non-nociceptive stimulation were selected: 25 – 65 ms (P50), 80 – 120 ms (P100), 100 – 220 ms (complex N140/P200), and 280 – 320 ms (P300). Similarly, the following time windows for nociceptive stimulation were selected: 100 – 160 ms (N100), 100 – 220 ms (N200), 300 – 500 ms (P300), 550 – 1000 ms (SCN).

The extracted ERPs time windows were subsequently statistically evaluated within 9 topographical regions of interest (ROIs; see Fig. 1 and 3): left frontocentral (FC1, FC3, FC5, TF7), right frontocentral (FC2, FC4, FC6, TF8), left central (C1, C3, C5, T7), right central (C2, C4, C6, T8), left centroparietal (CP1, CP3, CP5, TP7), right centroparietal (CP2, CP4, CP6, TP8), and the midline FCz, Cz, and CPz electrodes. Latencies and amplitudes of each ERP component provided in Table 1 were extracted at their dominant scalp electrode.

**Figure 1.**
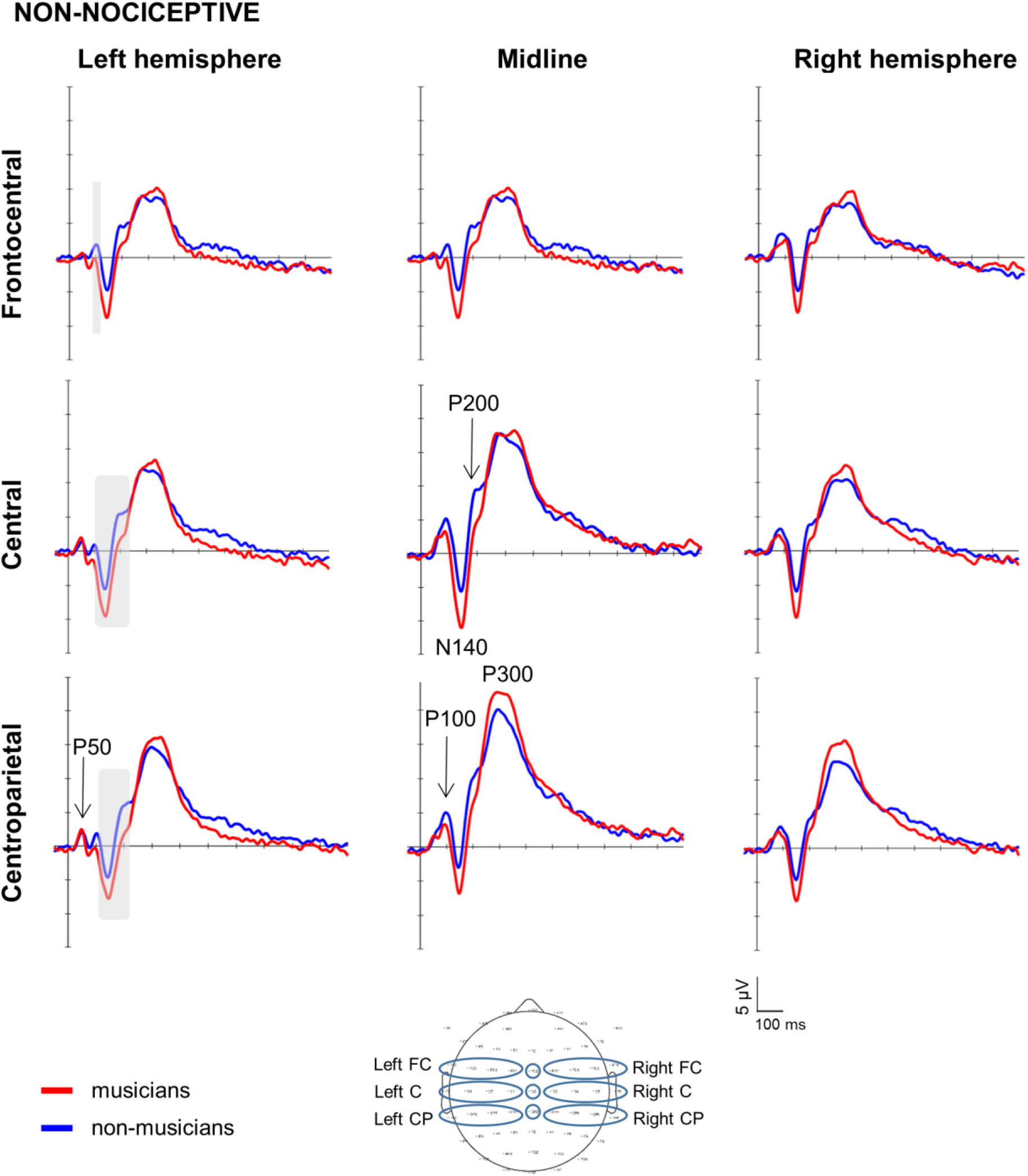
Brain responses to non-nociceptive somatosensory stimuli. Grand-averaged event related potentials elicited by the non-nociceptive electrical stimulation at the hand and illustrated at nine topographical regions of interest (ROIs; bottom center) in musicians (red lines) and non-musicians (blue lines). ROIs: frontocentral (FC), central (C), and centroparietal (CP). Marked time periods windows in grey indicate time periods and ROIs in which there were significant differences between musicians and non-musicians (p<0.05). Negative is plotted downward. Peak amplitudes across all ROIs were larger for the N140 and smaller for the P100 and P200 components in musicians compared to non-musicians.

**Figure 2.**
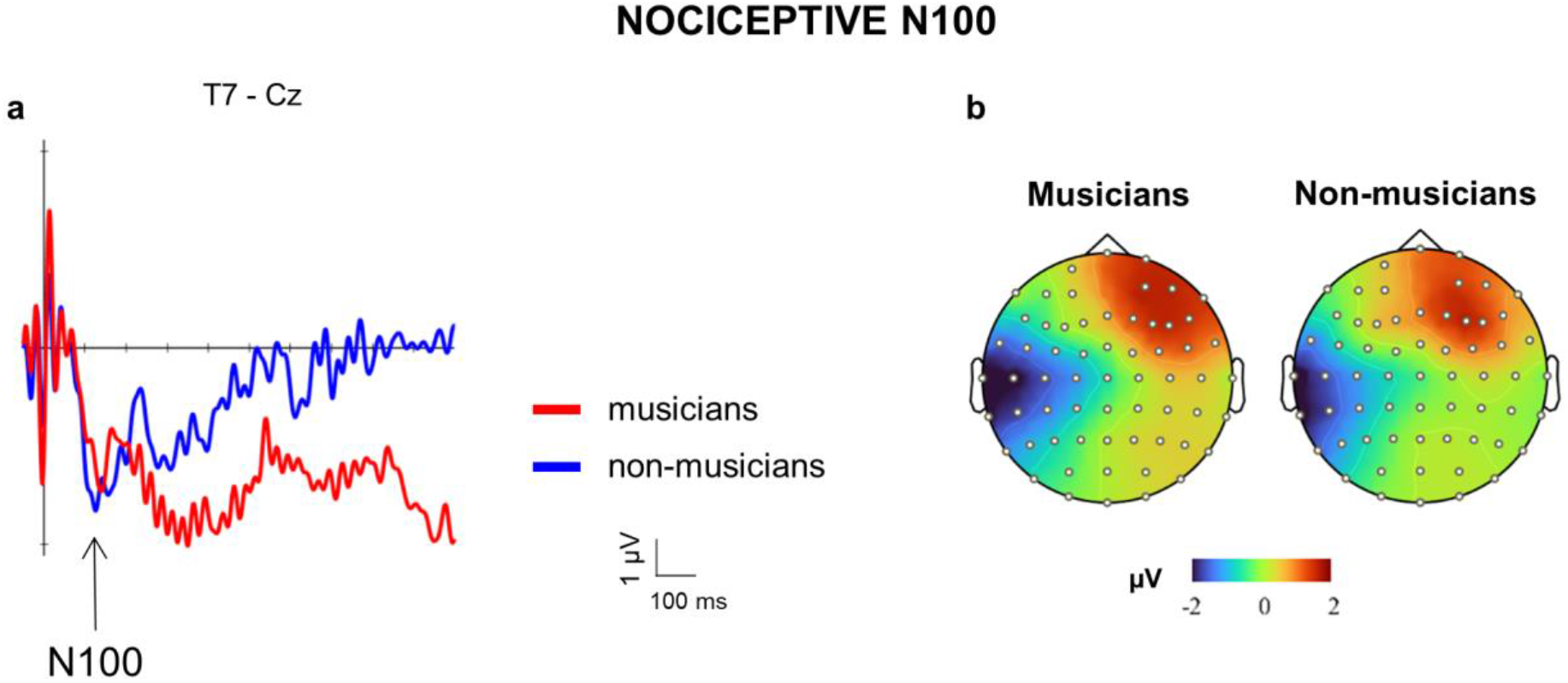
Brain responses and topography of nociceptive N100 component. **a)** Grand-averaged N100 event related potentials elicited by the nociceptive electrical stimulation at the hand and measured with the recommended temporal–frontal montage (T7-Fz) contralateral to the stimulated side (right hand). Musicians are represented by red lines and non-musicians by blue lines. Peak amplitudes of the N100 component indicated no significant differences between groups (p>0.5). **b)** Amplitude (in μV) scalp topography showing the distribution between musicians (left) and non-musicians (right) at 135 ms.

**Figure 3.**
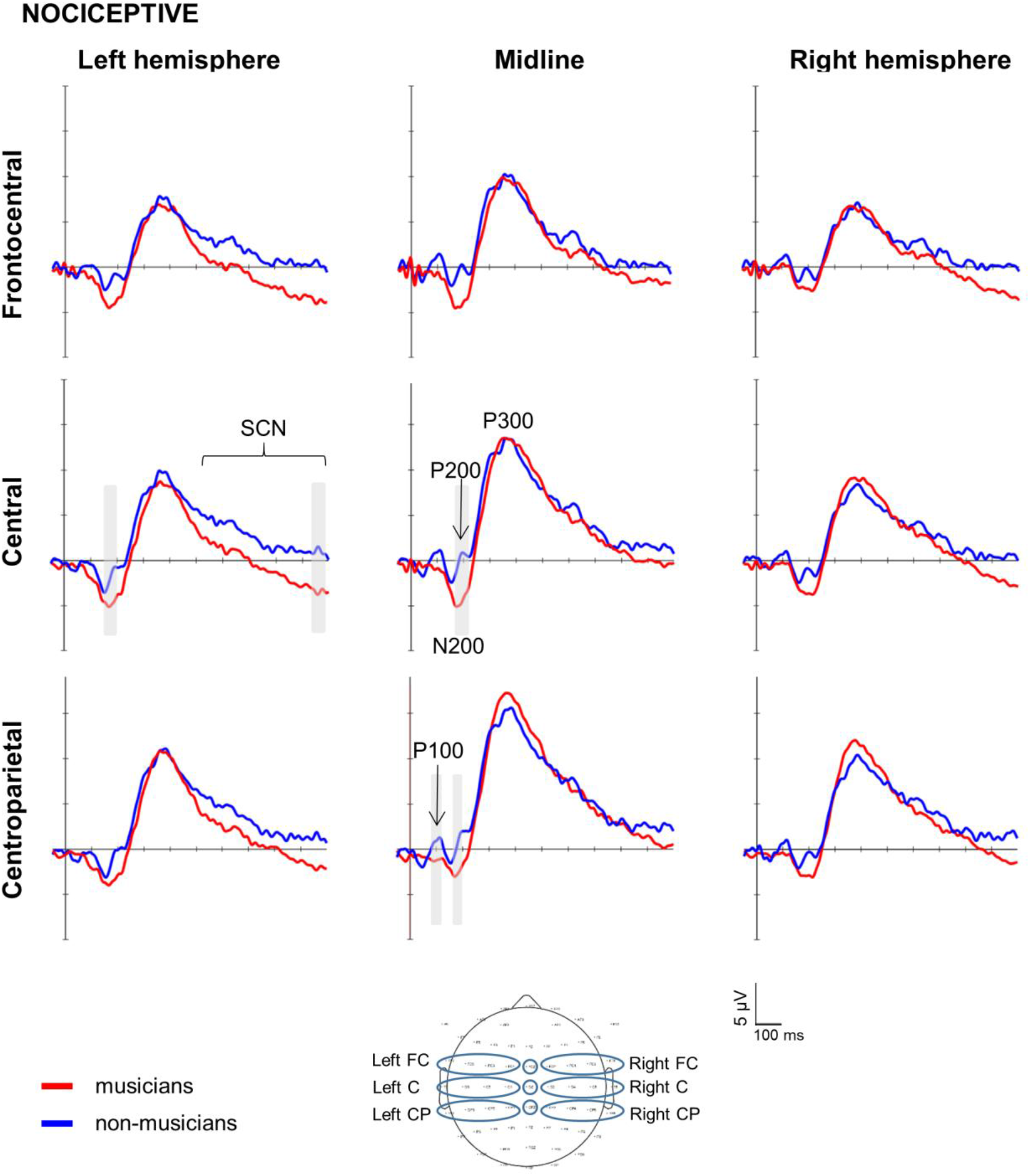
Brain responses to nociceptive stimuli. Grand-averaged event related potentials elicited by the nociceptive electrical stimulation at the hand and illustrated at nine topographical regions of interest (ROIs; bottom center) in musicians (red lines) and non-musicians (blue lines). ROIs: frontocentral (FC), central (C), and centroparietal (CP). Marked time periods windows in grey indicate time periods and ROIs with significant differences between musicians and non-musicians (p<0.05). Negative is plotted downward. Peak amplitudes across all ROIs indicate larger N200 and smaller P100 and P200 components, as well as a sustained contralateral negativity (SCN) in musicians compared to non-musicians.

### Statistical analysis

Data are presented in text and table as means and standard deviations. Behavioral responses were statistically analyzed with SPSS for Windows (IBM SPSS Statistics 26; IBM, Armonk, NY) and screened for assumptions of normality, sphericity, and homogeneity using descriptive plots and the Shapiro-Wilk’s, Mauchly’s, and Levene’s statistical tests. Detection thresholds for nociceptive and non-nociceptive stimuli were compared between groups using independent *t*-tests. Reaction times were analyzed using repeated measures analysis of variance (rmANOVA) with *Stimulation-modality* (nociceptive or non-nociceptive) as a repeated measures factor and *Group* (musicians and non-musicians) as a between group factor. Significant factors or interactions were analyzed post-hoc using Bonferroni’s procedure to correct for multiple comparisons.

ERPs were statistically analyzed with Brainstorm (Tadel et al., 2019, 2011), a freely available software released under the GNU general public license (http://neuroimage.usc.edu/brainstorm). ERP amplitudes and latencies in response to nociceptive and non-nociceptive stimuli were compared between groups using non-parametric permutation tests repeated 1000 times. False Discovery Rate (FDR) correction of multiple comparisons was employed to control for Type I errors (Benjamini and Hochberg, 1995).

Correlations and linear regressions were computed to investigate whether the ERPs to non-nociceptive and nociceptive stimulation could predict their respective stimulus detection thresholds and reaction times across all participants. In musicians, it was furthermore tested if accumulated musical training and daily practice time affected the amplitude and latency values of cortical ERPs, the reaction times, and the stimulus detection thresholds. A subset of electrophysiological responses was selected based on a priori hypotheses and correlation matrices. Specifically, we selected the amplitudes and latencies registered at the respective dominant scalp electrode (Cz) of the main biphasic nociceptive N200/P300 (N2/P2) components and their non-nociceptive N140/P200 equivalents. The amplitudes and latencies that showed no significant correlations with the dependent variables (i.e., reaction time and stimulus intensity) were excluded from the model. For all tests used, the level for significance was set at *p* < 0.05.

## RESULTS

### Behavioral measures

#### Stimulus characteristics

Nociceptive intra-epidermal electrical stimulation induced a pricking sensation in all participants, except for one subject of the non-musician group, who neither detected nor perceived the stimuli and was therefore excluded from the correponding analysis. Non-nociceptive electrical stimulation elicited a sensation of touch in all participants. Detection thresholds and reaction times confirmed that nociceptive and non-nociceptive stimulation selectively activated their corresponding fibers. That is, according to the kind of fibers and the physical characteristics of each electrode (Poulsen et al., 2020), nociceptive stimulation required lower stimulus intensity and generated slower reaction-times, in line with the characteristics of Aδ-fibers, whereas non-nociceptive stimulation required higher stimulus intensity and generated faster reaction-times, in line with the characteristics of Aβ-fibers.

#### Stimulus detection thresholds

No significant differences in detection thresholds were found for nociceptive (*t*_(31.08)_ = 1.51, *p* = 0.136) and non-nociceptive stimuli (*t*_(38)_ = −0.28, *p* = 0.777) between groups (Table 1).

#### Reaction times

Reaction time analysis (Table 1) revealed significant effects of *Stimulation-modality* (*F*_(1,37)_=47.014, *p* < 0.001) and *Group* (*F*_(1,37)_=7.198, *p* = 0.011). Post-hoc comparisons showed that reaction times were slower for the nociceptive than for the non-nociceptive stimulation in both groups *(p* < 0.001), and that musicians responded faster than non-musicians to both nociceptive and non-nociceptive stimuli (*p* < 0.05). The interaction between *Stimulation-modality* and *Group* was not significant (*F*_(1, 37)_=0.73, *p* = 0.398).

**Table 1.** Mean (± *SD*) detection thresholds, detection rates, reaction times, and cortical responses following nociceptive and non-nociceptive electrical stimulation. Peak latencies and amplitudes of each ERP component elicited by nociceptive (IEE) and non-nociceptive (ES) electric stimulation were respectively extracted at their dominant scalp electrode: nociceptive N1 (T7-Fz), nociceptive N2/P2 (Cz-A1); non-nociceptive P50 (CP3-A1); non-nociceptive N140/P200 (Cz-A1); non-nociceptive P100 and P300 (CPz-A1). ║: Significantly different within kind of stimulations (*p* < 0.001). *: Significantly different between groups (*p* < 0.05).

| **Intensity and Reaction times** | **Musicians**<br>**(n = 20)** | **Non-musicians**<br>**(n = 20)** |
| --- | --- | --- |
| Nociceptive detection threshold (μA) | 153.1 ± 63.5 | 192.5 ± 95.6 |
| Non-nociceptive detection threshold (μA) | 975.5 ± 316.6 | 948.5 ± 282.1 |
| Nociceptive detection rate (%) | 98 ± 5 | 92 ± 15 |
| Non-nociceptive detection rate (%) | 100 ± 1 | 98 ± 4 |
| Nociceptive reaction time (ms) | 354 ± 55* † | 433 ± 139* † |
| Non-nociceptive reaction time (ms) | 265 ± 48* † | 315 ± 89* † |
| **Latency (ms)** |  |  |
| *Nociceptive evoked potentials* |  |  |
| N100 | 136 ± 38 | 136 ± 39 |
| N200 | 182 ± 35 | 173 ± 35 |
| P300 | 330 ± 39 | 344 ± 44 |
| *Non-nociceptive evoked potentials* |  |  |
| P50 | 48 ± 4 | 48 ± 5 |
| P100 | 98 ± 9 | 100 ± 12 |
| N140 | 141 ± 10 | 141 ± 13 |
| P200 | 197 ± 11 | 192 ± 11 |
| P300 | 307 ± 26 | 312 ± 33 |
| **Amplitude (μV)** |  |  |
| *Nociceptive evoked potentials* |  |  |
| N100 | -7.79 ± 3.9 | -6.75 ± 2.7 |
| N200 | -9.32 ± 5.9 | -5.27 ± 4.9 |
| P300 | 15.68 ± 7.1 | 15.83 ± 6.4 |
| *Non-nociceptive evoked potentials* |  |  |
| P50 | 3.31 ± 2.9 | 2.75 ± 1.9 |
| P100 | 1.38 ± 5.6 | 3.77 ± 3.9 |
| N140 | -12.7 ± 10.5 | -7.25 ± 7.1 |
| P200 | 6.18 ± 7.2 | 9.72 ± 7.5 |
| P300 | 26.22 ± 6.9 | 20.9 ± 5.2 |

### Event-related potentials in response to non-nociceptive stimulation

All non-nociceptive ERP components were clearly identified (Fig. 1) in all participants except for one non-musician, in whom the EEG recording failed.

#### N140 – P200 analysis

The N140 – P200 complex exhibited a clear negative–positive biphasic wave with a maximum scalp-dominance at left (contralateral) central and midline Cz electrodes (Fig. 1 and 4). Visual inspection of peak amplitudes across all ROIs indicated a general enlargement of N140 and a reduction for the P200 component in musicians compared to non-musicians.

**Figure 4.**
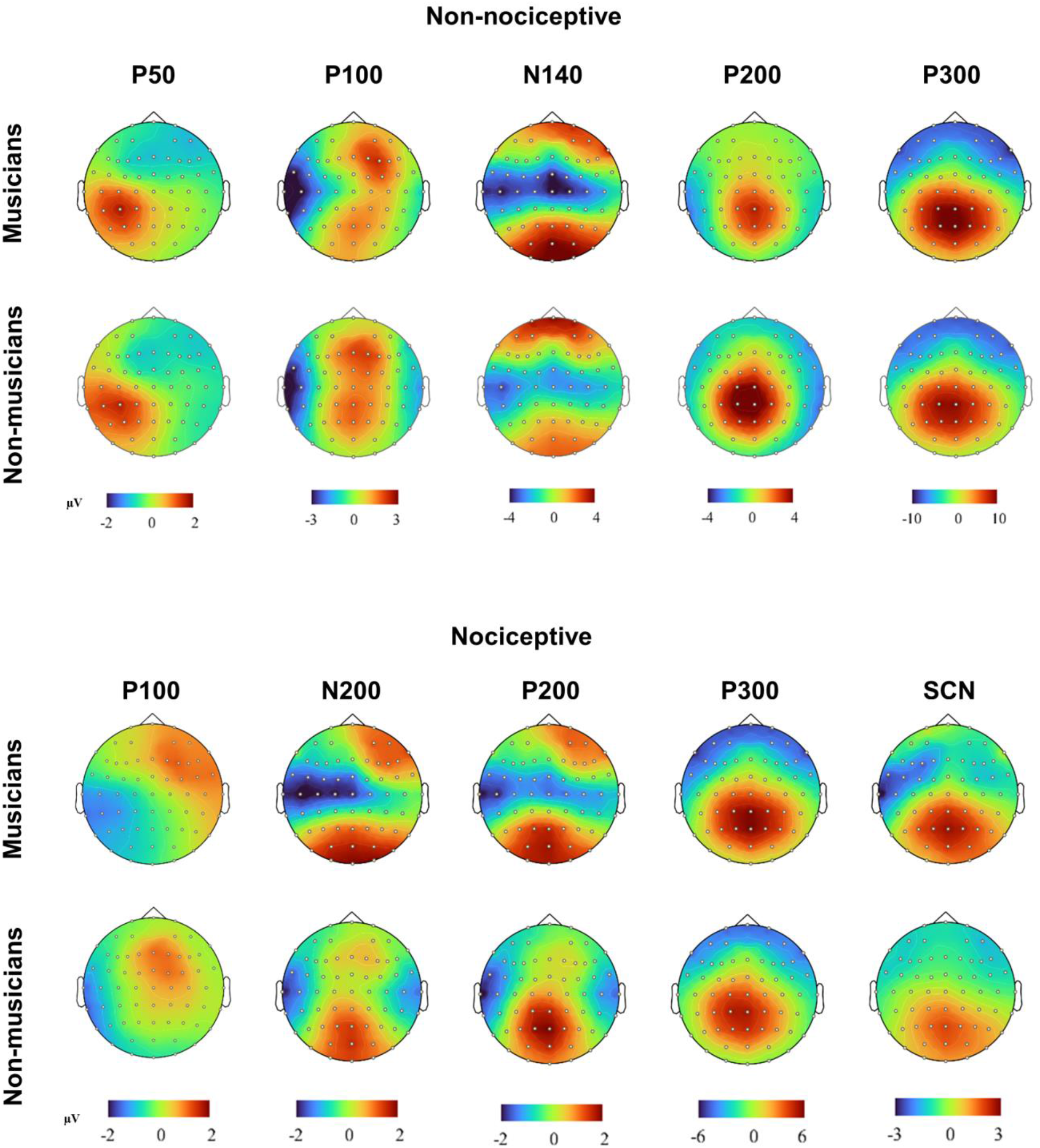
Scalp topographies to non-nociceptive and nociceptive stimuli. **Top:** Amplitude scalp topography of each non-nociceptive component in musicians and non-musicians. Scalp topographies shown are generated at 45 ms (P50); 100 ms (P100); 140 ms (N140); 200 ms (P200); and 300 ms (P300). **Bottom:** Amplitude scalp topography of each nociceptive component in musicians and non-musicians. Scalp topographies shown are generated at 100 ms (P100); 180 ms (N200); 200 ms (P200); 360 ms (P300), and 900 ms (SCN).

Peak amplitudes were significantly larger for the N140 and smaller for the P200 component in musicians compared to non-musicians at the contralateral (left) central (N140: *t*_(37)_ = 2.9, *p* < 0.05; P200: *t*_(37)_ = 2.7, *p < 0.05*) and centroparietal ROIs (N140: *t*_(37)_ = 2.8, *p < 0.05;* P200: *t*_(37)_ = 3.2, *p < 0.05*; Fig. 1).

Peak latencies extracted at Cz showed no significant group differences for the N140 or the P200 components at their dominant scalp electrodes (N140: *t*_(37)_ = 0.08, *p* = 0.93; P200: *t*_(37)_ = −1.05, *p* = 0.30; Table 1).

#### P50, P100 and P300 exploratory analyses

The exploratory analysis for P50 showed a left centroparietal-dominant scalp distribution contralateral to the stimulation side (45–55 ms after stimulus onset). The P100 peak amplitude scalp distribution (90–110 ms after stimulus onset) was frontocentral and centroparietal, whereas the P300 peak amplitude was distributed centroparietal and parietal (Fig. 1 and 4).

Peak amplitudes of the P50 and P300 components showed no significant group differences across the nine ROIs (all *t*_(37)_ < 1.44; *p* > 0.15; Fig. 1 and 4). However, peak amplitudes for the P100 time window yielded a significantly smaller P100 amplitude in musicians compared to non-musicians for the left frontocentral ROI (*t*_(37)_ = 3.1; *p* < 0.05; Fig. 1).

Peak latencies of the P50 extracted at CP3-A1, as well as P100 and P300 extracted at CPz-A1 showed no significant group differences (all *t*_(37)_ < 0.48; *p* > 0.53; Table 1).

### Event-related potentials in response to nociceptive stimulation

#### N100

The N100 component was clearly identified in all non-musicians and in 19 out of 20 musicians (Fig. 2). Peak amplitudes of the N100 component, measured with the recommended temporal–frontal montage (T7-Fz), revealed no significant group differences (*t*_(36)_= 0.948; *p >* 0.5; Table 1). Peak latencies of the N100 component revealed no significant group differences either (*t*_(36)_= 0.948; *p* > 0.05; Table 1).

#### N200 – P300 complex

Nociceptive stimulation elicited a clear vertex potential constituted by a negative–positive biphasic wave (N200 – P300 complex) in all participants with a scalp dominance at central midline electrode (Fig. 3 and 4). Visual inspection indicated enlarged peak N200 amplitudes (Fig. 3) in musicians compared to non-musicians. In addition, visual inspection of the N200 component time window also indicated a prominent positive component at 200 ms (labelled P200) in non-musicians but not in musicians.

Peak amplitudes of the N200 component were larger in musicians compared to non-musicians at the left central (*t*_(37)_ = 3.2, *p* < 0.05), midline central (*t*_(37)_ = 3.5, *p* < 0.05), and midline centroparietal ROIs (*t*_(37)_ = 3.1, *p* < 0.05). Moreover, nociceptive P200 peak amplitude was smaller in musicians compared to non-musicians at the left and midline central (*t*_(37)_ = 2.9, *p* < 0.05 and *t* = 3.7, *p* < 0.05, respectively) and at the centroparietal midline ROIs (*t*_(37)_ = 2.7, *p* < 0.05).

Peak amplitudes of the P300 component revealed no significant group differences across ROIs (all *t*_(37)_ < 1.5; *p* > 0.63). Peak latencies of the N200 and P300 components revealed no significant group differences at their dominant scalp electrodes (all *t*_(37)_ < 0.73; *p* > 0.32; Table 1).

Visual inspection of peak amplitudes also indicated a prominent component at 100 ms (labelled P100) at dominant frontal and central midline scalp distributions in the non-musicians group (Fig. 3 and 4). The peak P100 amplitude was significantly smaller in musicians compared to non-musicians at the midline centroparietal ROI (*t*_(37)_ = 3.0, *p* < 0.05). Peak latencies of the P100 component revealed no significant group differences (*t*_(37)_ = −0.483, *p* = 0.63).

### Sustained contralateral negativity

Following the main nociceptive components, a gradually developing SCN in musicians compared to non-musicians was observed (Fig. 3 and 4). This SCN component was visually larger in musicians compared to non-musicians. The topographical maps in Figure 4 illustrate the scalp distribution of SCN, indicating a focus over left (contralateral), frontocentral and central ROIs. The SCN amplitude was bigger in musicians compared to non-musicians at the left (contralateral) central ROI (*t*_(37)_ = 2.4, *p* < 0.05; Fig. 3).

### Across participants, non-nociceptive and nociceptive components correlate with reaction times

Reaction times showed a positive significant correlation with their respective non-nociceptive N140 (*r* = 0.49, *p* = 0.002) and nociceptive N200 (*r* = 0.35, *p* = 0.029) peak amplitudes (Fig. 5A), but not with their respective latencies. Reaction times did not significantly correlate with the non-nociceptive P200 and nociceptive P300 latencies and amplitudes (all *p* > 0.05).

**Figure 5.**
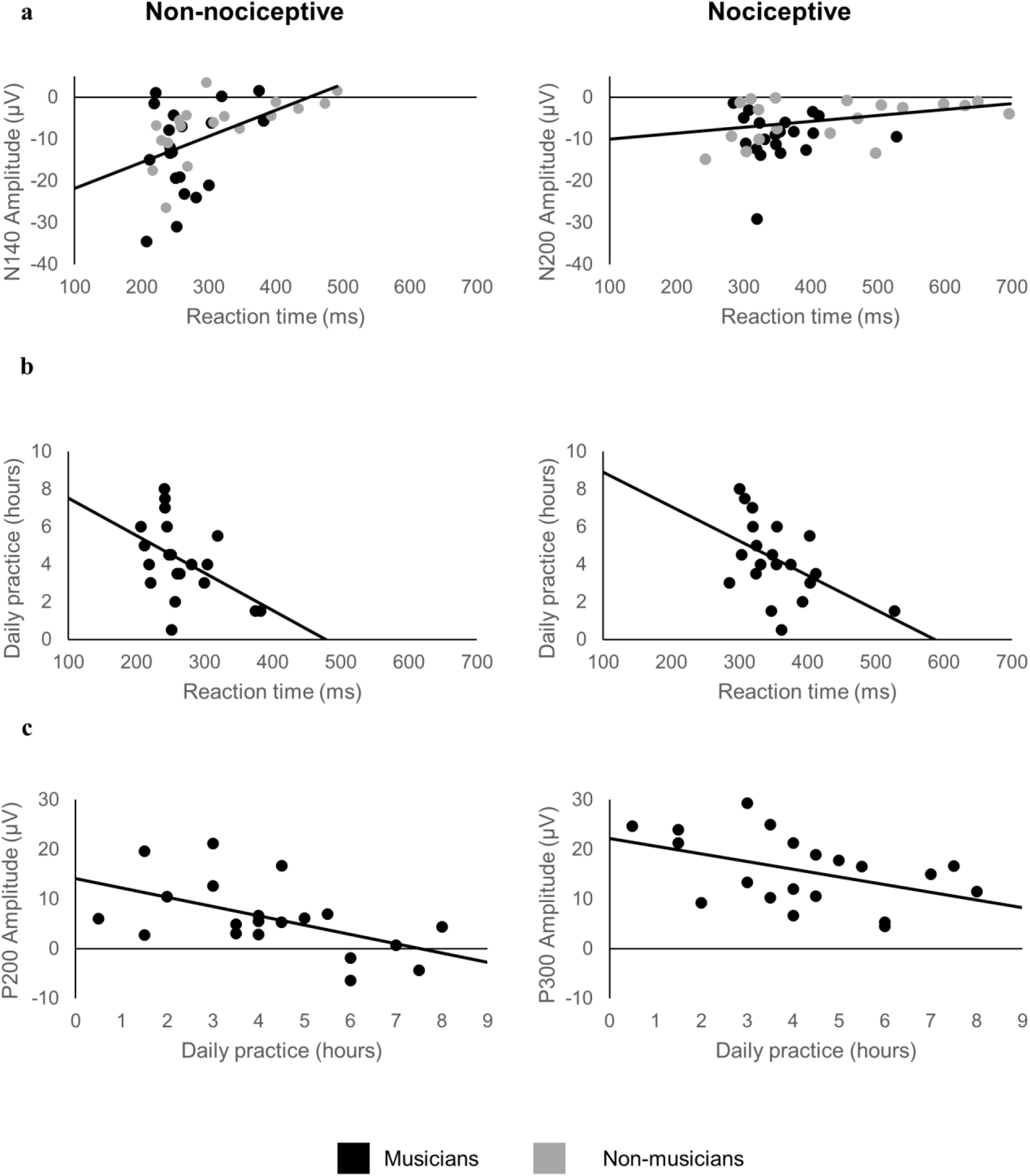
Significant correlations of event related potentials, reaction times, and daily practice. **a)** The non-nociceptive N140 (left) and nociceptive N200 (right) peak amplitudes correlate with their respective non-nociceptive and nociceptive electrical stimulation reaction times. Musicians are represented by black dots and non-musicians by grey dots. **b)** In musicians, the amount of daily practice (h) correlates with the peak amplitudes of the non-nociceptive P200 and the nociceptive P300 components. **c)** In musicians, the amount of hours of daily practice correlates with the reaction times of the non-nociceptive and the nociceptive electrical stimulation. Fit lines indicate correlations between respective variables.

### Across participants, the non-nociceptive P200 and nociceptive P300 correlate with stimulus detection thresholds

Stimulus detection thresholds for the non-nociceptive stimulus showed a negative significant correlation with the magnitude of the corresponding P200 peak amplitudes (*r* = −0.40, *p* = 0.010). Non-nociceptive detection thresholds were neither significantly correlated with the P200 latency nor with the N140 amplitude and latency (all *p* > 0.05).

Stimulus detection thresholds for the nociceptive stimulus showed a negative significant correlation with the latencies of the corresponding P300 peak (*r* = −0.43, *p* = 0.006). Nociceptive detection thresholds were not significantly related with the respective nociceptive N200/P300 amplitudes nor with the latencies of the N200 (all *p* > 0.05).

### The amount of daily sensorimotor practice in musicians correlates with non-nociceptive and nociceptive components as well as reaction times

In musicians, the amount of daily practice showed a negative significant correlation with the non-nociceptive (*r* = −0.47, *p* = 0.035) and nociceptive (*r* = −0.49, *p* = 0.027) reaction times (Fig. 5B), as well as with the magnitude of the non-nociceptive P200 (*r* = −0.53, *p* = 0.016) and the nociceptive P300 (*r* = −0.45, *p* = 0.048) amplitudes (Fig. 5C).

In order to test the relationship of daily practice on the magnitude of the evoked response amplitudes, we included musicians in a regression, using stimulus intensity and daily practice as predictors. Higher non-nociceptive detection thresholds predicted smaller P200 amplitudes, accounting for 22.7% of the variance (F_(1,19)_ = 5.29, *R^2^* = 0.23, *p* = 0.034). By adding daily musical practice, the model explained 43.8% of the variance (*F*_(2, 19)_ = 6.61, *R^2^* = 0.44, *p* = 0.008). Thus, more daily practice significantly improved the prediction of smaller P200 amplitudes (*F*_*change* (1,19)_ = 6.36; *R^2^_change_* = 0.21, *p* = 0.022), accounting for an additional 21.1% of the variance.

## DISCUSSION

Using musicians as a model for use-dependent plasticity, the present study investigated whether prolonged and repeated sensorimotor training and its associated multisensory integration may alter the neural mechanisms of pain processing in healthy individuals. We observed distinct differences in transient brain responses and behavioral reaction times to nociceptive and non-nociceptive somatosensory stimuli as a function of sensorimotor training. Specifically, healthy musicians showed larger N200 peak amplitudes to nociceptive stimulation as well as larger N140 and smaller P200 peak amplitudes to non-nociceptive stimulation, and displayed faster RTs. Notably, daily sensorimotor training in musicians predicted RTs, non-nociceptive P200, and nociceptive P300 amplitudes, emphasizing the use-dependent nature of this modulation. Moreover, larger non-nociceptive N140 and nociceptive N200 components predicted faster RTs across all participants. ERP waveforms to nociceptive stimulation were characterized by a gradually developing sustained contralateral negativity in musicians, which did not appear during non-nociceptive stimulation. This novel evidence provides first direct support for a putative model, proposing that the same mechanisms by which repetitive sensorimotor training and multimodal integration can enhance selectivity to non-nociceptive stimuli may also facilitate neural responses to nociceptive cues in healthy humans.

### Effects of extensive sensorimotor training on non-nociceptive processing

Behavioral and electrophysiological effects of sensorimotor musical training on non-nociceptive stimulus processing have been well documented. Specifically, within superior acoustical processing of fine-grained timbre, rhythm, and pitch discrimination in musicians (Kraus and Chandrasekaran, 2010; Vuust et al., 2008) and correspondingly enlarged auditory evoked potentials following music-dependent specialization (Fujioka et al., 2006; Pantev et al., 1998). In addition, accumulated sensorimotor training has also been linked with enhanced tactile acuity and enlarged cortical representation of the digits in the somatosensory cortex (Elbert et al., 1995; Ragert et al., 2004).

In the present study, extensive sensorimotor training facilitated upstream perceptual information processing and top-down response control (reaction time) during non-nociceptive electrical stimulation, which was used to assess the effects of sensorimotor training on the dorsal column-lemniscal pathway, as indicated by enlarged N140 and decreased P200 amplitudes. Across all participants, the N140 amplitude predicted the reaction times to stimulus detection, analogous to previous reports (Talsma et al., 2007), whereas in musicians, the hours of daily musical training predicted both the shorter reaction times and the decreased P200 amplitudes. Considering that sensorimotor experience in musicians has been linked to enhanced multimodal perception and correspondingly faster motor responses (Landry and Champoux, 2017), the current results suggest that the N140/P200 complex may reflect the electrophysiological correlate of this superior behavioral performance.

In the general population, enhanced N140 components have been reported in response to multisensory (tactile and visual) relative to unisensory stimulation (Eimer and Driver, 2000; Ohara et al., 2006), supporting the theory that multimodal integration facilitates a more robust sensory perception that shapes cross-modal effects on evoked neural responses (Driver and Noesselt, 2008; Talsma et al., 2010). As outlined above, musical training has long been known to enhance auditory evoked potentials such as the N100 (Pantev et al., 1998; Shahin et al., 2004), a negative peak recorded between 100–200 ms that is equivalent to the tactile N140 and the nociceptive N200. Similar effects have been reported in non-musicians after one year of musical training (Fujioka et al., 2006), indicating that the temporal synchrony of neurons may be augmented through musical experience. Also, the early somatosensory cortical responses are enhanced in trumpet players compared to non-musicians when tactile (lip stimulation) and auditory (trumpet tones) cues are presented simultaneously (Schulz et al., 2003), thus providing evidence of cross-modal reorganization associated with multimodal sensorimotor training. The nature of this cortical facilitation is moreover task-specific, as enhanced N100 potentials in musicians have been linked to the timbre of their principal instrument (Pantev et al., 2001).

Within a musical framework, skilled performance requires the precisely timed integration, segregation, and prediction of ongoing auditory, visual, tactile, proprioceptive, and visceral feedback (Kleber et al., 2013; Lee and Noppeney, 2011; Schirmer-Mokwa et al., 2015), which has also been associated with neural adaptations related to enhanced (insula-based) salience detection and attentional selectivity, as well as increased functional connectivity between the insula, cingulate, and somatosensory cortices to facilitate the access to the motor system (Kleber et al., 2017; Luo et al., 2014; Schirmer-Mokwa et al., 2015; Zamorano et al., 2017). Taking this evidence into account, the present data suggest that extensive sensorimotor training and corresponding multisensory integration may specifically facilitate the priming of neural responses in brain areas where multimodal stimuli converge (Lu et al., 2014; Murray et al., 2005; Paraskevopoulos et al., 2012).

### Effects of extensive sensorimotor training on nociceptive processing

Larger N200 amplitudes were found in the current study during nociceptive stimulation in healthy musicians compared to healthy non-musicians. Moreover, the individual N200 amplitudes across all participants predicted the reaction times to nociceptive stimulation, where reaction times were generally faster in musicians. As the N200-P300 components are predominantly generated in the insula, frontal operculum, and cingulate cortex (Garcia-Larrea et al., 2003), these results may indicate higher temporal synchrony and neuronal firing among these regions as a function of musical training. This notion is supported by previous resting-state fMRI studies, in which healthy musicians compared to healthy non-musicians showed increased temporal correlation in blood oxygenation level dependent (BOLD) signals between the insular cortex and the cingulate, orbitofrontal, and dorsolateral prefrontal cortices (Zamorano et al., 2017). In addition, these studies found comparably enhanced insula-based connectivity in healthy musicians and in chronic pain non-musicians (Zamorano et al., 2019), suggesting that extensive sensorimotor training and chronic pain may trigger adaptations in overlapping neural systems. The larger nociceptive N200 amplitudes in the current study also support previous behavioral findings of increased pain sensitivity in healthy professional musicians (Zamorano et al., 2015). Together, data from the current study provide the first direct evidence in healthy human individuals that supports the hypothesis that extensive and repetitive sensorimotor training can modulate the neural mechanisms underlying the processing of pain.

Increased electrophysiological activity in insular and cingulate cortices has also been observed in experimental pain models of secondary hyperalgesia. These studies showed that short periods of sustained nociceptive input delivered by high-frequency nociceptive stimulation (HFS) on the skin not only induce hypersensitivity and faster reaction times, but also enhance the N200 ERP components elicited by activation of A-δ and C-fiber nociceptors around the stimulated area (Biurrun Manresa et al., 2018; Lenoir et al., 2018). The underlying mechanisms following sustained nociceptive HFS have been associated with long-term potentiation (LTP) of excitatory synaptic transmission, a key feature for improving signal processing and sensory transmission (Froemke et al., 2013), between peripheral nociceptive fibers within dorsal horn neurons projecting to the parabrachial area in the brainstem (Ikeda et al., 2003). In addition, sustained nociceptive HFS can also enhance the non-nociceptive evoked N100 peak amplitude (the negative peak recorded between 100–200 ms and equivalent to N140 and N200) in response to vibrotactile (van den Broeke and Mouraux, 2014) and visual stimulation (Torta et al., 2018, 2017), suggesting that sustained nociceptive stimulation may also engage LTP mechanisms in supra-spinal multisensory areas, such as the insula and ACC (Zhuo, 2014).

Sustained high frequency non-nociceptive stimulation (i.e., transcutaneous electrical nerve stimulation, TENS), on the other hand, diminishes pain perception, leads to hypoalgesia, and induces a reduction of the N100, N200, and P300 amplitudes to nociceptive stimuli (Peng et al., 2019), contrary to the effects of accumulated extensive sensorimotor training on nociceptive pathways reported in the current study. An explanation for the different findings between experimental (TENS) and the ecological model of non-nociceptive stimulation used in the current study (i.e. musical training) may be a difference in their underlying neural mechanisms. That is, experimental non-nociceptive TENS is a passive unisensory stimulation performed for only 20 min (Sluka and Walsh, 2003), activating Aβ fibers and inhibiting incoming nociceptive inputs transmitted via Aδ and C fibers at the spinal level with the contribution of a supraspinal descending inhibitory mechanism (Peng et al., 2019). Musical training, in contrast, represents long-term (i.e., years of) active multisensory stimulation and motor integration, which is known to facilitate supra-spinal LTP-like mechanisms (Bütefisch et al., 2000; Zatorre et al., 2013) that may be similar to the mechanisms of nociceptive HFS. Thus, LTP-like mechanisms associated with musical training may not only facilitate the transmission of task-specific non-nociceptive sensory inputs, but possibly also enhance multisensory signal processing at the spinal and supra-spinal pathways, which facilitate the perceptual processing of nociceptive signals.

### Effects of extensive sensorimotor training on sustained contralateral negativity after nociceptive stimulation

In an attempt to explore additional effects of extensive sensorimotor training beyond the targeted components, a prominent sustained negativity was identified in musicians contralateral to the stimulated hand. This effect only occurred at contralateral electrodes in response to nociceptive stimulation in musicians starting after 300 ms post-stimulus and persisting throughout the inter-stimulus interval. The sustained negativity has previously been attributed to the maintenance of sensory information in working memory and corresponding behavioral differences in working memory capacity (Katus et al., 2015; Vogel et al., 2005; Vogel and Machizawa, 2004). Although the current experiment did not challenge working memory, the similar characteristics suggest that the perceptual processing of nociceptive inputs in musicians may have activated a selective retention of noxious inputs in working memory. However, the exact neural basis of this effect needs to be determined in future work.

### Extensive sensorimotor training and neural adaptations in nociceptive pathways

Based on the present data, we propose a tentative explanatory model in which extensive sensorimotor learning and the associated multisensory integration may not only trigger adaptive task-specific neuroplasticity and correspondingly enhanced behavioral performance, as commonly reported in the use-dependent plasticity literature (Elbert et al., 1995; Herholz and Zatorre, 2012), but also induces supra-spinal LTP-like mechanisms that may simultaneously facilitate the processing of nociceptive and non-nociceptive somatosensory signals in brain areas where multimodal inputs converge (i.e., the insula). Such a neurobiological mechanism may plausibly explain the observed increase in stimulus-selectivity to nociceptive cues in healthy individuals performing repetitive movements, as demonstrated in our musician model (Zamorano et al., 2015). This explanation is furthermore supported by studies demonstrating that multisensory integration leads to a more robust percept (Ernst and Bülthoff, 2004), induces cross-modal plasticity in multisensory conversion zones (Driver and Noesselt, 2008), and facilitates nociceptive neural and behavioral responses, as indicated by invertebrate models (Hu et al., 2017; Ohyama et al., 2015).

### Conclusion

In the current study, we described the cortical mechanisms that link extensive sensorimotor training and corresponding multisensory integration, both known triggers of use-dependent plasticity, to neural adaptations in nociceptive pathways using experienced musicians as an ecologically model. Enhanced neural responses to electrical nociceptive and non-nociceptive somatosensory stimulation in musicians relative to non-musicians provides the first direct evidence for a link between altered processing of nociceptive inputs and repetitive sensorimotor training in healthy humans. These novel findings may contribute to the understanding of the high variability in neural responses to nociceptive stimulation in the general population and extend current putative models that explain the increased vulnerability for exacerbated pain sensitivity prevalently found in individuals performing repetitive movements. Further neurophysiological research using experimental models of persistent pain is warranted to investigate if these neural adaptations may be considered a risk factor for developing chronic pain.

